# Platelet factor 4 modulates endothelial cell antimicrobial activity to enhance bacterial clearance and improve sepsis outcomes

**DOI:** 10.1101/2025.09.02.673783

**Authors:** Anh T. P. Ngo, Weronika Ortmann, Abigail Skidmore, Hyunjun Kim, Jenna Oberg, Amrita Sarkar, Veronica Bochenek, Nate Levine, Lubica Rauova, Irina Chernysh, Zachary Martinez, Caroline Diorio, Mark Goulian, Victor Nizet, Mortimer Poncz, Kandace Gollomp

## Abstract

Sepsis is a life-threatening condition characterized by dysregulated host responses to infection. Here, we identify platelet factor 4 (PF4) as a key mediator of vascular antimicrobial defense. In vitro, PF4 enhanced endothelial cell internalization of *Escherichia coli* via interactions with the PF4 receptor CXCR3 and the endothelial glycocalyx, directing bacteria to clathrin-mediated endocytosis and lysosomal degradation. In vivo, PF4 administration improved survival and reduced sepsis severity, bacterial burden, inflammation, and thrombosis in wild-type (WT) and PF4 knockout (*PF4*^−*/*−^) mice challenged with systemic polymicrobial infection. Using intravital microscopy, we observed that infused bacteria were rapidly sequestered in the pulmonary microvasculature. However, *PF4*^−*/*−^ mice exhibited impaired bacterial clearance and increased microvascular platelet adhesion and aggregation. In the liver, following Kupffer cell depletion, *PF4*^−*/*−^ mice had increased sinusoidal platelet accumulation, larger bacterial aggregates, and elevated hepatic bacterial burden compared to WT controls. Collectively, these findings reveal that PF4 promotes bacterial clearance and restrains immunothrombosis during sepsis in part via endothelial cell uptake and destruction of microbes. By enhancing endothelial antimicrobial function, PF4 represents a significant yet previously underrecognized host defense mechanism that limits bacterial spread and alleviates vascular injury during infection.

**KEY POINTS:**

- In vitro, PF4 accelerates bacterial clearance by enhancing endothelial uptake of bacteria and promoting their trafficking to the lysosome.
- In murine sepsis, PF4 augments pathogen clearance to reduce infection severity, limit organ injury, and improve survival.

## INTRODUCTION

Sepsis is a life-threatening condition caused by a dysregulated immune response to infection that remains a leading cause of mortality worldwide. Current treatments for sepsis-induced multiorgan dysfunction are limited to antibiotics and supportive care.^1^ Beyond their traditional role in hemostasis and thrombosis, platelets have emerged as key mediators of inflammation and innate immunity.^2^ Thrombocytopenia, a common feature of sepsis, is associated with increased morbidity and mortality,^3^ underscoring the importance of platelets in host defense. Platelets contribute to antimicrobial responses by recognizing pathogen-associated molecular patterns (PAMPs), activating leukocytes, and releasing immunomodulatory mediators.^4,5^

One such mediator is platelet factor 4 (PF4; CXCL4), a member of the CXC chemokine family and the most abundant protein in platelet α-granules.^4^ Upon activation, platelets release PF4 at concentrations exceeding 12 μg/mL.^6^ High level of expression across most mammalian species suggest an evolutionarily conserved function.^6^ Our group has previously demonstrated that PF4- deficient (*PF4*^*-/-*^) mice exhibit increased mortality in both lipopolysaccharide (LPS)-induced endotoxemia and the cecal ligation and puncture (CLP) model of polymicrobial sepsis.^7,8^ Furthermore, exogenous PF4 treatment mitigates sepsis severity, reduces markers of thrombosis, and improves survival, supporting a protective role for PF4 in infection.^7,8^

Unlike other CXC chemokines, PF4 lacks the N-terminal Glu-Leu-Arg (ELR) motif required for activation of leukocytes via G-protein-coupled chemokine receptors CXCR1 and CXCR2.^9,10^ Instead, PF4 forms cationic tetramers that bind polyanions with high affinity, a property central to many of its proposed biological functions.^11^ Many of PF4’s effects arise from electrostatic interactions with anionic partners, including glycosaminoglycans (GAGs) on leukocytes,^12^ polyphosphates,^13^ von Willebrand factor,^14^ and nucleic acids.^15^ PF4 also binds to anionic components of bacterial cell walls, such as LPS on gram-negative bacteria and teichoic acid on gram-positive organisms, forming antigenic complexes that elicit anti-PF4/polyanion antibodies.^16,17^ These antibodies opsonize PF4-coated bacteria, enhancing their phagocytosis.^16,17^ PF4 also binds to the negatively-charged DNA scaffold of neutrophil extracellular traps (NETs), promoting NET resistance to nuclease digestion while enhancing bacterial trapping and reducing NET-mediated thrombogenicity.^8,18^

The vascular endothelium is another major binding partner for PF4. It is coated by an anionic glycocalyx, composed of heparan sulfate-rich proteoglycans and glycoproteins, that interact with PF4 with high affinity,^12^ though less strongly than heparin.^19^ PF4 tetramers also engage the CXCR3B receptor expressed on endothelial cells (ECs),^20^ positioning PF4 as an effective opsonin capable of bridging bacteria to the endothelium. With an estimated 1-6×10^13^ ECs covering approximately 4,000 m^2^ of surface area, the endothelium plays a crucial frontline role in sepsis by protecting underlying tissue from blood-borne pathogens and toxins.^21^ Beyond their barrier function, ECs are among the first cells to detect and respond to invading bacteria. They express pattern recognition receptors (PAMPs) and damage-associated molecular patterns (DAMPs),^22^ as well as antimicrobial peptides such as β-defensin 3, RNase 7, and cathelicidin LL-37.^23^ Accumulating evidence indicates that ECs contribute to immune defense by clearing apoptotic cells, cell debris, and microbes, and by recruiting immune cells to vulnerable vascular sites to limit pathogen spread.^24–27^ However, the intrinsic antimicrobial pathways of ECs, potentially including phagocytosis, autophagy, and innate immune signaling, remain incompletely understood.

In this study, we investigated the mechanisms by which PF4 influences bacterial clearance during infection. To achieve this, we developed in vitro models demonstrating that PF4 enhances bacterial internalization by cultured ECs, followed by lysosomal trafficking and degradation. In a systemic polymicrobial bacteremia model, PF4-opsonized bacteria were cleared more efficiently in vivo, with reduced systemic inflammation and thrombosis. These findings suggest that PF4 facilitates a previously unrecognized mechanism of pathogen recognition and clearance, contributing to host defense during sepsis.

## MATERIALS and METHODS

All reagents were purchased from Sigma-Aldrich except as noted. Recombinant human PF4 was expressed in S2 cells and purified using affinity chromatography and protein liquid chromatography as previously described.^44^ The end-product was found to be endotoxin-free using ToxinSensorTM chromogenic LAL endotoxin assay kit (Genscript; L00350) and was tested for size distribution by SDS-polyacrylamide gel electrophoresis.

### Scanning electron microscopy

Live *E. coli* (1×10^8^ CFU/ml) were incubated with PF4 (0-100µg/ml) and then plated on 10mm glass slides. samples were fixed in 2% of glutaraldehyde 0.1M cacodylate buffer with 0.15 M Na Cl, pH 7.4, dehydrated with ethanol started from 30% up to 100% and then finally with 100% hexamethyldisilazane (Acros Organics, Fair Lawn, NJ), and sputter-coated with gold-palladium as described by Vorjohann et al.^45^ High-definition micrographs were obtained by FEI Quanta 250FEG scanning electron microscope (FEI, Hillsboro, OR) from multiple representative areas of each sample. Fiji (NIH) open-source image processing software was used to quantify bacterial cluster size.

### Endothelial cell bacterial uptake and killing studies

Human umbilical vein endothelial cells (HUVECs; Lifeline Cell Technology, FC-0044) were grown to confluence on 0.1% gelatin–coated (Stem Cell Technologies; 07903) glass-bottom 8- well Ibidi chambers (Ibidi; 80826) with Vasculife VEGF endothelial medium complete kit (Lifeline; LL-0003). In select experiments, HUVECs were pre-treated with TNF⍰(20 ng/mL; R&D Systems; 210-TA) for 30 min at 37°C to induce inflammation prior to bacteria exposure. Quiescent or inflamed HUVECs were incubated with heat-inactivated *S. aureus*-pHrodo (2×10^6^/mL; Invitrogen; P35367) in serum-free media (SFM; Gibco; 11-111-044) or together with PF4 (1.25-100 μg/mL) overnight at 37°C. To assess *S. aureus* bioparticle uptake, HUVECs were then washed with phosphate-buffered saline (PBS; Gibco; 10010049) and imaged using Zeiss 10× and 20× objectives on a Zeiss LSM 710 confocal microscope. Mean fluorescence intensity (MFI) was quantified using Fiji. All MFI values were normalized to MFI of cells that did not receive PF4 treatment.

To measure VWF release following overnight *S. aureus* bioparticle (2×10^6^/mL) exposure, HUVECs were washed with PBS and fixed in 4% paraformaldehyde (Santa Cruz Biotechnology; sc-281692) for 15 min at RT before blocking with 3% bovine serum albumin (A2153) and 2% fetal bovine serum (Cytiva; SH30071.03) in PBS for 1 hr at room temperature (RT). Primary anti–human VWF antibody (1:1000 dilution in PBS; Agilent Dako, A0082) was incubated overnight at 4°C. Slides were washed with PBS and blocked for 1 hr at RT. Secondary Alexa Fluor 594 anti–rabbit IgG (4 μg/mL; Invitrogen, A32740) and Hoechst 33342 (10 μg/mL; Invitrogen, H3570) were incubated in PBS for 2 hr at RT in the dark. HUVECs were then subjected to confocal microscopy, and MFI was quantified using Fiji.

HUVECs were grown to confluence on 0.1% gelatin-coated 24-well plates and pre-treated with TNF⍰(20 ng/mL). In select experiments, inflamed HUVECs were pre-treated with AMG-487 (20 µg/mL; MedChemExpress; HY-15319), Dynasore (100 µM; Tokyo Chemical Industry; D5461),

Pitstop (30 µM; Abcam; AB120687), mixture of heparinases I, II, and III (1 U/mL; BBI Solutions), or hydroxychloroquine (HCQ; 50 µM; Selleckchem; S4430) for 45 min at 37°C prior to bacterial exposure. *E. coli* (K-12 strain) was grown in Luria Bertani (LB) broth with shaking at 200 rpm at 37°C until reaching log-growth phase (OD_600_ = 0.4; ∼3×10^8^ colony forming units (CFUs) per mL). Inflamed HUVECs were then exposed to live *E. coli* at multiplicity of infection (MOI) of 10 in SFM alone or in combination with PF4 (2.5 and 25 μg/mL) for 1 hour at 37°C. ECs were then washed with PBS to remove unbound bacteria and incubated for up to 5 hr at 37°C.

### Lactate dehydrogenase assay

HUVEC culture supernatant was collected every hour to quantify lactate dehydrogenase (LDH) release as a measure of cytotoxicity (Promega; J2380). Cells were washed with PBS and treated with 1% penicillin-streptomycin (Gibco; 15140122) in SFM for 1 hr at 37°C to kill extracellular bacteria. Cells were then washed with PBS and lysed by gentle scraping. Lysates were diluted in PBS and plated on LB agar plates for 24 hr at 37°C for CFU enumeration.

### Cecal slurry systemic polymicrobial sepsis model

Cecal contents from at least 3 wildtype (WT) C57BL/6J mice (Jackson Laboratory, strain 000664). were resuspended in 5% dextrose (cecal slurry; CS) and stored at −80°C at concentrations of 400 mg/mL. Twelve-to-16-week-old WT and *PF4*^*-/-*^ mice on a C57BL/6J background received an intravenous (I.V) injection of CS (20 mg) or CS (20 mg) pre-incubated with PF4 (100 μg) at 37°C for 20 min. Sepsis severity (MSS, body temperature) and mortality were assessed up to 96 hr post CS challenge. Blood was drawn from the retro-orbital sinus into EDTA 1 week before CS challenge (baseline), at 6 hr, and at 24 hr for hematological analyses, CFU enumeration, and platelet-poor plasma isolation by centrifugation at 2,000 × g for 20 min at RT.

To determine bacterial burden 24 hr post infection, a subset of WT mice received an I.V injection of CS (80 mg) or CS (80 mg) pre-incubated with PF4 (100 μg) at 37°C for 20 min. Since 80 mg CS killed all *PF4*^*-/-*^ mice before 24 hr, a subset of *PF4*^*-/-*^ mice received an I.V injection of CS (20 mg) or CS (20 mg) pre-incubated with PF4 (100 μg) at 37°C for 20 min. 24 hr post infection, blood and spleen, lung, and liver homogenates were subjected to CFU enumeration.

### Proximity Extension assay

Plasma protein profile was analyzed by Proximity Extension Assay technology (Olink Proteomics, Uppsala, Sweden) and included 92 proteins from the Olink Target 96 Mouse Exploratory panel related to inflammation, metabolism, cell development, tissue injury, and immune regulation. Relative protein concentration was expressed as NPX (log_2_ normalized protein expression). To assess thrombin generation in septic plasma, thrombin anti-thrombin (TAT) complexes were measured by ELISA (AssayPro; EMT1020-1) according to manufacturer’s instructions. Briefly, EDTA-plasma was diluted 110-fold and incubated for 2 hr at RT in 96-well plates pre-coated with polyclonal antibody against mouse thrombin. The wells were washed and incubated with biotinylated mouse TAT complex antibody for 1 hr at RT. The microplate was then incubated with streptavidin-peroxidase conjugate for 30 min at RT, and chromogenic substrate was added for 5 min at RT. Reactions were stopped and absorbance was immediately measured at 450 nm. Von Willebrand factor (VWF) levels were measured in the ETDA-plasma from CS-challenged mice by ELISA (Abcam; ab208980) according to manufacturer’s instructions. Briefly, plasma samples were diluted 110-fold and incubated with anti-mouse VWF A2 capture and detector (HRP-conjugated) antibodies for 1 hr at RT in 96-well plates pre-coated with an anti-tag antibody. The wells were washed, and TMB development solution was added for 10 min at RT in the dark. Reactions were stopped and absorbance was immediately measured at 450 nm.

### In situ-microscopy studies

#### Lung intravital microscopy

Intravital confocal microscopy was used to visualize *E. coli*, platelets, and neutrophils in the lungs of 8–12-week-old WT and *PF4*^*-/-*^ mice following bacteria injection. Mice were anesthetized with intraperitoneal (I.P) injection of Nembutal (11 mg/kg) and a tracheal cannula was inserted and attached to a MiniVent ventilator (Havard Apparatus). Mice were ventilated with a tidal volume of 10 μL of compressed air per gram of mouse weight, a respiratory rate of 130-140 breaths per minute, and a positive-end expiratory pressure of 2 cm H_2_O. The mice were placed in the right lateral decubitus position and the surface of the left lung was exposed. A thoracic window was applied and secured to the stage. Twenty-five mm Hg of suction was applied using a vacuum regulator (Amvex) to immobilize the lung. The confocal microscope 50× objective was lowered into place over the thoracic window. Mice were pre-infused with anti-CD41 and anti- Ly6G antibodies to detect platelets and neutrophils, respectively. Animals then received jugular infusion of GFP-expressing *E. coli* (1×10^9^ CFU) in 200 μL of PBS administered over 20-30 seconds. Images were analyzed using Imaris (Oxford Instruments), SlideBook (3i), and ImageJ (NIH).

#### Liver intravital microscopy

Mice were anesthetized with Nembutal (11 mg/kg), followed by cannulation of the right jugular vein for administration of fluorescent antibodies and GFP-expressing *E. coli*. Liver preparation for intravital imaging was performed as previously described.^46^ Briefly, with the animal in a supine position, a skin incision was made in the lower abdominal quadrant and extended along the midline until the level of the sternum. To expose the liver, a second incision was made along the linea alba at the center of the peritoneal membrane, extending from the lower abdomen to the xiphoid cartilage. After liver exposure, the ligament connecting the gallbladder to the diaphragm was cut. A glass coverslip was placed over the exposed left liver lobe prior to immersion of the objective lens. Imaging was performed using an Olympus BX61WI microscope (Olympus, Tokyo, Japan) with a 20×/0.50 numeric aperture (NA) water-immersion objective lens.

#### Depletion of liver Kupffer cells (KCs)

To deplete KCs, mice received I.P. injections containing clodronate liposomes (Liposoma, Amsterdam, The Netherlands; 10 µL per 1 g body weight) 3 days prior to intravital imaging per the manufacturer’s protocol.^34^ Control mice received an I.P. injection containing PBS loaded liposomes. Successful KC depletion was confirmed by F4/80 antibody infusion and liver intravital imaging.

### Statistics

Data are presented as mean ± standard error of the mean (SE). The Shapiro-Wilk normality test was used to determine whether group data were distributed normally. Levene’s test was used to determine equality of variances. One-way ANOVA with Tukey’s post hoc test was used to compare between treatment groups. Mann-Whitney test or Kruskal-Wallis with Dunn’s post hoc test was used to compare between groups when the data did not qualify for parametric statistics. A *P* value of 0.05 or less was considered significant. All statistical analyses were conducted using GraphPad Prism 9.

## RESULTS

### In vitro characterization of PF4-bacterial interactions

Previous studies have shown that PF4 binds to surface polyanions on both gram-positive and gram-negative bacteria, enhancing their phagocytosis by macrophages and neutrophils.^16,17,28^ To further characterize PF4-bacterial interactions, we used scanning electron microscopy (SEM) to examine the association of PF4 with live *Escherichia (E*.*) coli*. As shown in **Fig. 1A**, PF4 induced dose-dependent bacterial aggregation, increasing the average *E. coli* cluster size from 3.09 μm^2^ in the absence of PF4 to 4.06 μm^2^ and 5.24 μm^2^ with 20 and 100 μg/mL of PF4, respectively.

**Figure 1.**
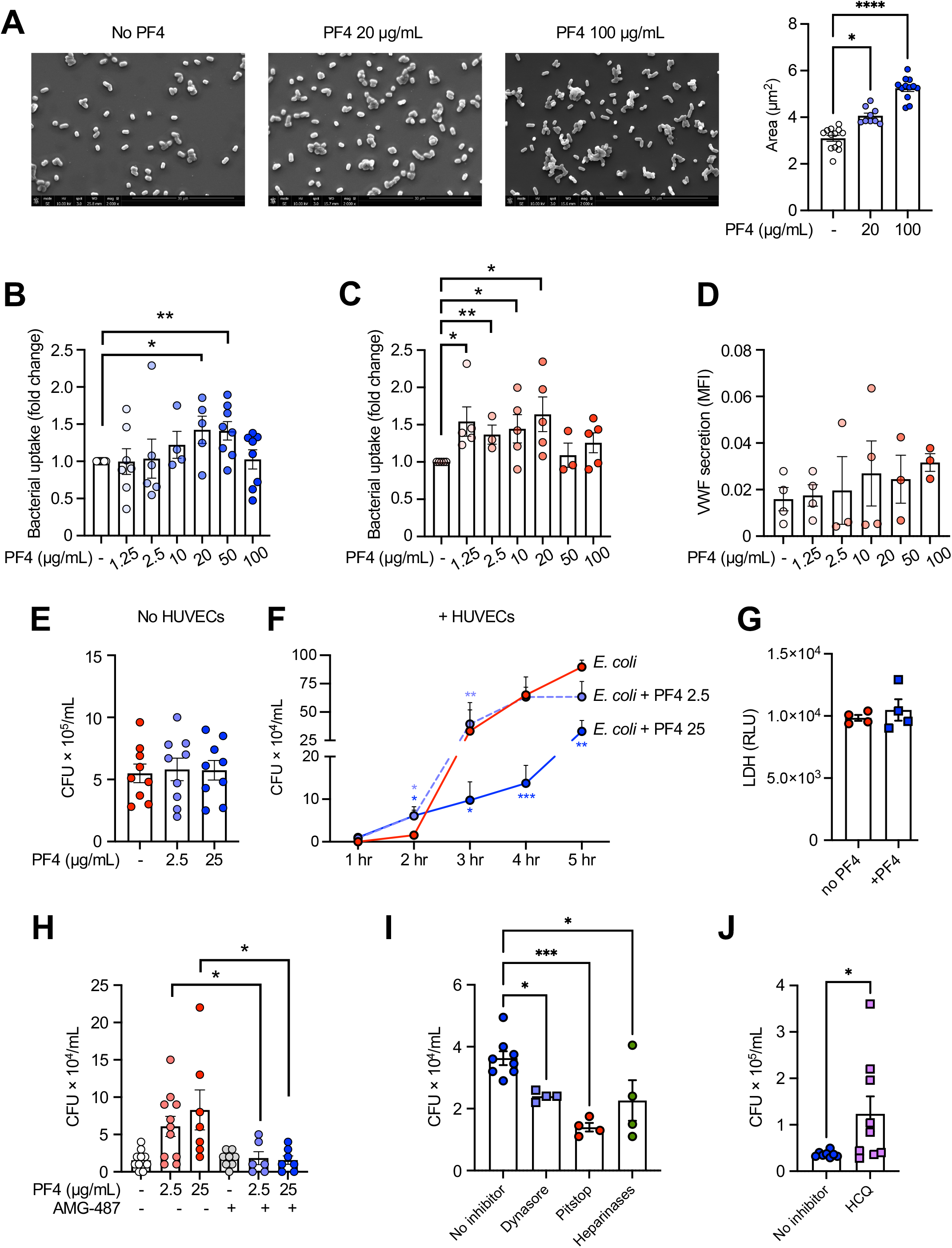
PF4 aggregates bacteria and enhances endothelial uptake and killing. (**A**) Representative SEM images and quantification of *E. coli* aggregate area ± PF4 (0-100 µg/mL). (**B**) HUVEC uptake of pHrodo conjugated *S. aureus* bioparticles (2×10^6^/mL) ± PF4 (0-100 µg/ml). (**C**) Same as (**B**) but following HUVEC treatment with TNF⍰(20 ng/mL) to model inflammation. MFI was normalized to untreated controls. (**D**) TNF⍰-treated HUVECs were exposed to *S. aureus* bioparticles pre-incubated with increasing PF4 concentrations and VWF release was quantified as a surrogate marker of endothelial activation. (**E**) *E. coli* was incubated with PF4 (0, 2.5, and 25 µg/mL) for 20 min at 37ºC and plated for CFU enumeration. (**F**) *E. coli* was pre-incubated with PF4 (0, 2.5, or 25 µg/mL) for 20 min at 37ºC prior to exposure to TNF⍰- treated HUVECs. At hourly intervals, cells were treated with antibiotics to kill extracellular *E. coli*, and CFU enumeration was performed on EC lysates to quantify internalized bacteria. (**G**) LDH release was measured in HUVEC supernatants after 2 hr incubation with *E. coli*. (**H**) TNF⍰-treated HUVECs were incubated with vehicle or AMG-487 (20 µg/mL) for 45 min prior to exposure to *E. coli* ± PF4 (2.5 or 25 µg/mL). Two hours post infection, ECs were washed, treated with antibiotics, and cell lysates were subjected to CFU enumeration. (**I**) TNF⍰-treated HUVECs were treated with Dynasore (100 µM), Pitstop (30 µM), or heparinases (1 U/mL), (**J**) or HCQ (50 µM) for 45 min prior to exposure to *E. coli* ± PF4 (25 µg/mL) at 37ºC. Two hours post infection, cells were treated with antibiotics, and lysates were subjected to CFU enumeration. Mean ± SE. ^*^*P* < 0.05, ^**^*P* < 0.01, ^***^*P* <0.001, ^****^*P* < 0.0001.

### PF4 enhances endothelial uptake of Staphylococcus (S.) aureus bioparticles

To assess whether PF4 promotes bacterial uptake by ECs, we incubated quiescent HUVECs with pHrodo-conjugated *S. aureus* bioparticles, which fluoresce upon internalization into acidic endosomal and lysosomal compartments, in the presence of increasing PF4 concentrations. Uptake followed a bell-shaped curve, rising at 20 and 50 μg/mL and declining at 100 μg/mL (**Fig. 1B**), similar to the pattern observed with PF4 complexing to heparin.^29^ To mimic the inflammatory environment of sepsis, HUVECs were pretreated with tumor necrosis factor (TNF) α (20 ng/mL) for 30 min. Under these conditions, the uptake curve shifted leftward, with PF4 now enhancing EC-bioparticle uptake at lower concentrations (1.25-20 μg/mL), while concentrations of 50 μg/mL and above had no effect (**Fig. 1C**).

To determine whether PF4-induced bacterial uptake leads to endothelial activation, we measured von Willebrand factor (VWF) release as a surrogate marker of EC reactivity. Inflamed HUVECs exposed to *S. aureus* bioparticles with increasing PF4 concentrations (as in **Fig. 1C**) did not exhibit a significant change in VWF staining, indicating that PF4-enhanced phagocytosis does not induce general endothelial activation (**Fig. 1D**).

### PF4 enhances endothelial uptake and intracellular killing of phagocytosed E. coli

To determine whether PF4 exerts direct antimicrobial activity, we incubated live *E. coli* with increasing PF4 concentrations and measured CFUs. No differences were observed between vehicle- and PF4-treated groups (**Fig. 1E**), consistent with prior studies showing that PF4 is not directly bactericidal.^17,30,31^

We next evaluated whether PF4 enhances bacterial clearance through endothelial uptake and intracellular killing. Inflamed HUVECs were exposed to PF4-pretreated *E. coli* (0-25 µg/mL) for 1 hr. Cultures were then washed and treated with antibiotics to eliminate extracellular bacteria. To measure subsequent growth of internalized bacteria, HUVECs were lysed at the indicated time points and CFUs were quantified. Two hours post-exposure, HUVECs incubated with bacteria treated with 25 μg/mL of PF4 exhibited significantly higher intracellular CFUs compared to untreated controls, demonstrating enhanced uptake (**Fig. 1F**). However, from 3 to 5 hr, intracellular CFUs declined in the 25 μg/mL PF4 group, falling below levels observed in untreated or a low dose of 2.5 μg/mL PF4-treated cells (**Fig. 1F**), suggesting that while PF4 promotes early bacterial uptake, it also accelerates intracellular bacterial clearance.

To evaluate whether PF4-induced *E. coli* uptake compromises host cell integrity, we measured lactate dehydrogenase (LDH) release in the supernatant 2 hr after EC infection. LDH levels did not differ between PF4-treated (25 μg/mL) and untreated HUVECs (**Fig. 1G**), suggesting that PF4-induced bacterial uptake is not cytotoxic to host cells.

### Characterization of PF4-bacteria-endothelial cell interaction

PF4 binds CXCR3B, an alternatively spliced isoform of the CXCR3 chemokine receptor expressed on ECs, which mediates angiostatic responses and is expressed on the endothelial cell surface.^20^ To evaluate whether this interaction contributes to bacterial uptake, Inflamed HUVECs were treated with the small-molecule CXCR3 inhibitor AMG-487 for 2 hr prior to incubation with PF4-opsonized *E. coli* (2.5 and 25 μg/mL). CXCR3 inhibition significantly reduced PF4-mediated bacterial internalization (**Fig. 1H**), implicating CXCR3B as a receptor in this process.

To further investigate the mechanism of uptake, we enzymatically removed endothelial surface heparin and heparan sulfate GAGs using heparinase treatment,^32^ which similarly reduced internalization of PF4-treated *E. coli* (**Fig. 1I**). Pharmacological blockade of receptor-mediated endocytosis with Dynasore (a dynamin inhibitor) and Pitstop (a clathrin-mediated endocytosis inhibitor) also decreased *E. coli* internalization (**Fig. 1I**), supporting that PF4-opsonized bacteria are internalized through clathrin-dependent pathways.

Finally, we assessed whether internalized bacteria are trafficked to lysosomes for degradation. Treatment of HUVECs with hydroxychloroquine (HCQ), which alkalinizes lysosomes and impairs their degradative capacity, resulted in a marked rise in intracellular *E. coli* (**Fig. 1J**). Together, these findings suggest that PF4 promotes bacterial uptake through interaction with CXCR3 and GAGs on the endothelial surface, leading to clathrin-mediated endocytosis and subsequent trafficking to lysosomes for intracellular killing.

### PF4 improves outcomes in a murine model of systemic polymicrobial sepsis

We next investigated whether PF4 treatment improves outcomes in vivo using a cecal slurry (CS) model of systemic polymicrobial sepsis. WT and *PF4*^*-/-*^ mice received tail vein infusion of CS (20 mg/mouse), either alone or pre-incubated with PF4 (100 μg/mouse; CS + PF4). Animals were then monitored for 96 hr. Within 15 min of CS infusion alone, all the mice developed hypothermia and exhibited a hunched posture, which resolved within 1 hr, while these effects were not seen in mice administered CS + PF4 (data not shown). Blood cultures obtained 6 hr after CS infusion showed no bacterial growth (data not shown). However, by 24 hr, clinical signs of illness returned, including hypothermia, elevated murine sepsis scores (MSS).^33^ *PF4*^*-/-*^ mice challenged with CS showed more rapid progression to mortality compared to WT mice (100% vs 33% mortality by 48 hr), although this difference did not reach statistical significance (**Fig. 2A**). Notably, all animals that received CS alone expired before 72 hr, whereas 100% of WT mice receiving CS+PF4 were protected from mortality, and 62.5% of *PF4*^*-/-*^ mice treated with CS+PF4 were protected (**Fig. 2A**). PF4 treatment also significantly reduced MSS and prevented hypothermia in WT mice at 6 hr, and in *PF4*^*-/-*^ mice 24 hr post-challenge (**Fig. 2B**).

**Figure 2.**
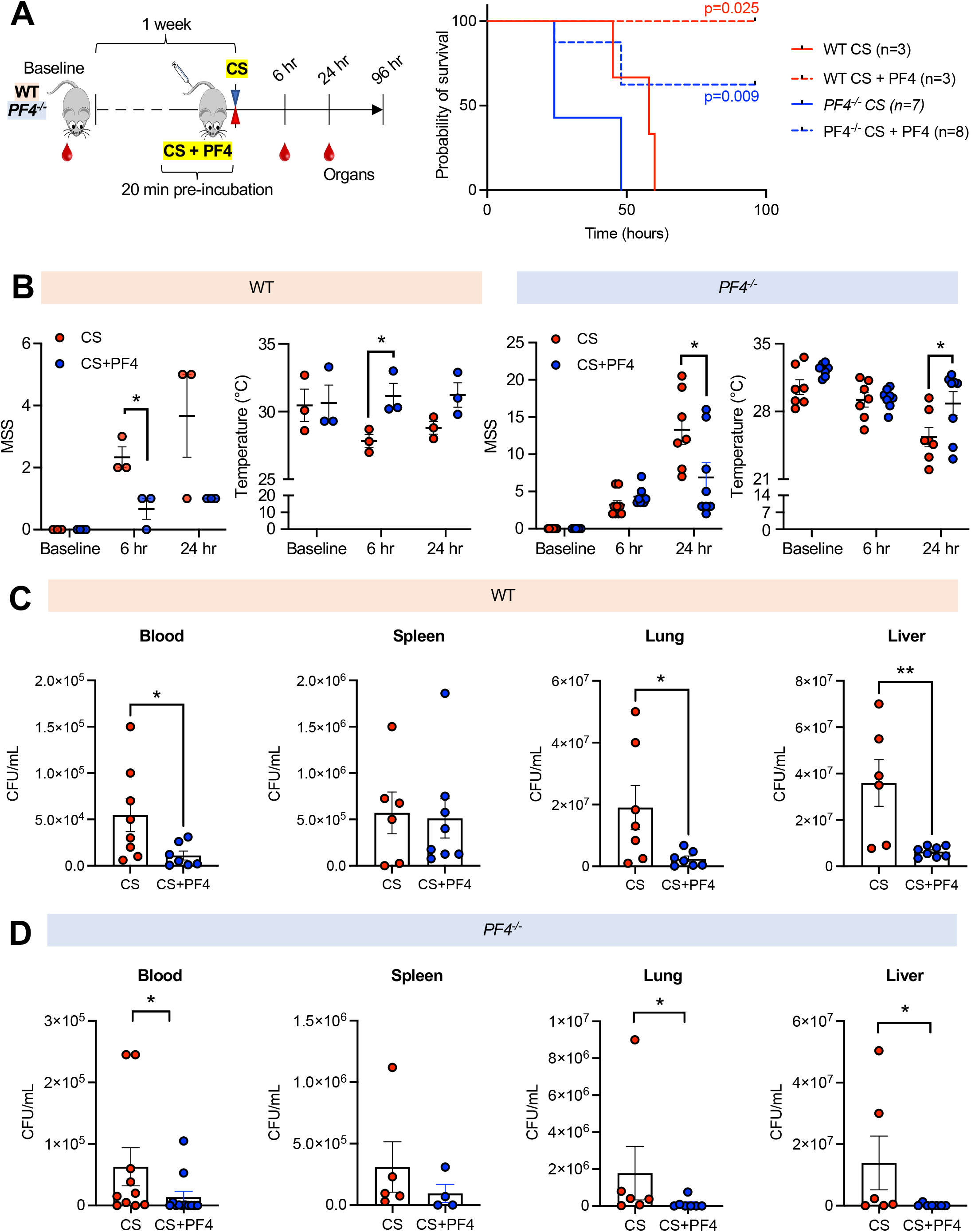
PF4 improves survival outcomes and promotes bacterial clearance in murine polymicrobial sepsis. (**A-B**) WT (red) and *PF4*^*-/-*^ (blue) mice received CS (20 mg/mouse; solid line), or CS + PF4 (100 µg PF4/mouse; dotted line) via tail vein injection. (**A**) Survival was assessed for up to 96 hr. (**B**) MSS and temperature were assessed 6 and 24 hr post CS challenge. (**C-D**) A separate cohort of WT (**C**) and *PF4*^*-/-*^ (**D**) mice received CS (80 mg CS per WT mouse; 20 mg CS per *PF4*^*-/-*^mouse), diluted blood and organ homogenates were subjected to CFU enumeration at 24 hr post infection. Mean ± SE. ^*^*P* < 0.05, ^**^*P* < 0.01.

Since blood cultures obtained at 6 hr post CS infusion at 20 mg/mouse dose showed no bacterial growth, we performed a separate set of experiments using a higher CS dose of 80 mg/mouse to assess systemic bacterial burden. Notably, all *PF4*^*-/-*^ mice receiving this dose died before 24 hr (data not shown), whereas WT mice survived. Accordingly, in subsequent experiments we compared bacterial load at 24 hr post-infection in WT mice challenged with 80 mg CS (**Fig. 2C**) and *PF4*^*-/-*^ mice challenged with 20 mg CS (**Fig. 2D**). PF4 co-treatment significantly reduced CFUs in blood, lung, and liver in both WT and *PF4*^*-/-*^ mice compared to genotype-matched mice given CS alone.

### PF4 reduces levels of inflammatory and pro-thrombotic markers in sepsis

To identify biological pathways modulated by PF4 during infection, we analyzed plasma from WT mice 24 hr after CS challenge using the Olink Target 96 Mouse Exploratory panel. CS- induced sepsis led to elevated levels of pro-inflammatory cytokines interleukin (IL) 6, IL10 and CSF2 (granulocyte-macrophage colony-stimulating factor; GM-CSF), as well as the chemokine CCL5 (RANTES) (**Fig. 3A**). Markers associated with tissue damage and remodeling, including cardiac troponin (TNNI3), integrin αvβ6 (ITBG6), TNF receptor superfamily member 12A (TNFRSF12A), and transforming growth factor beta receptor III (TGFBR3), were also increased (**Fig. 3A**). PF4 co-treatment significantly attenuated these inflammatory and injury-associated responses (**Fig. 3A**).

**Figure 3.**
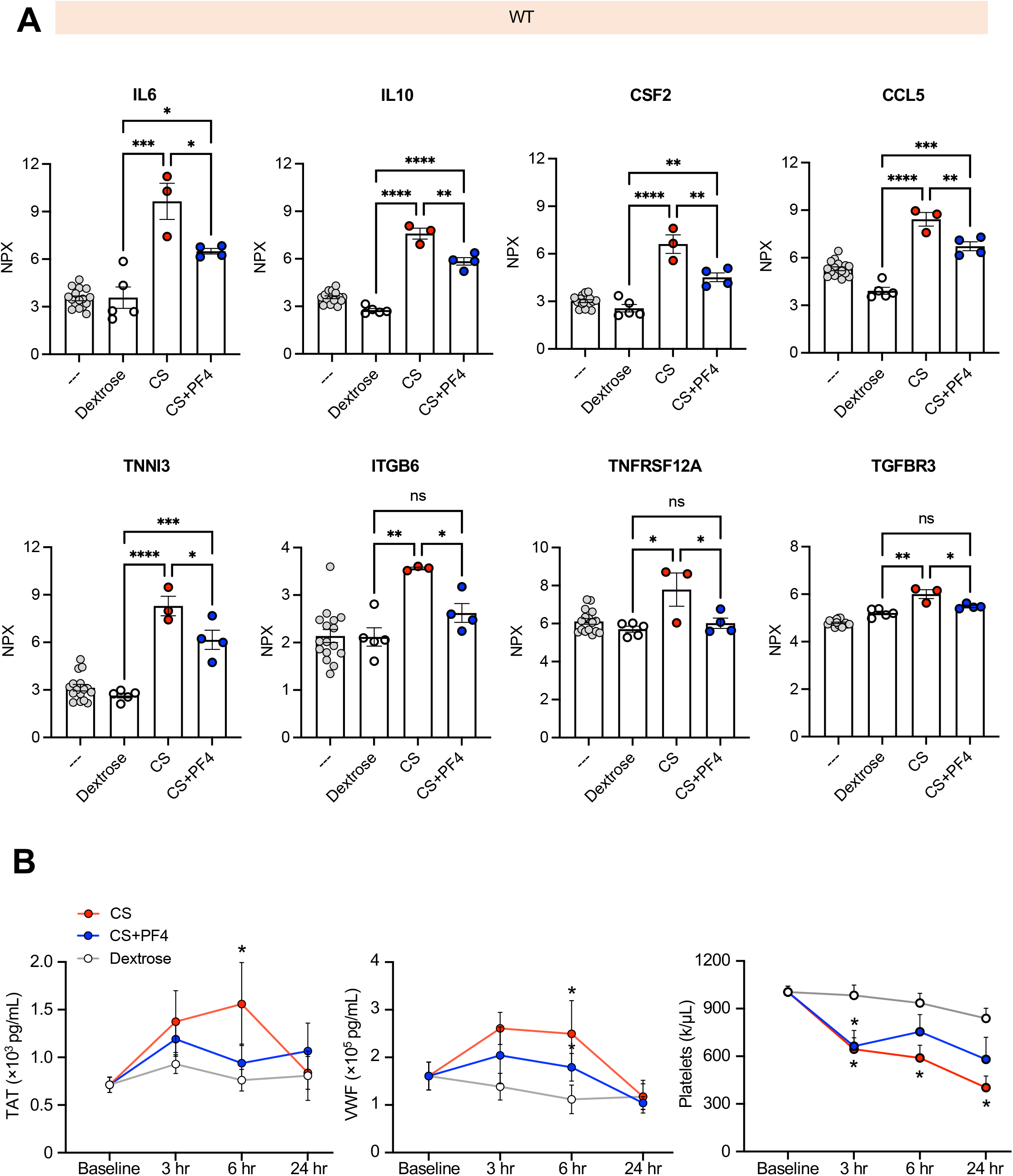
Effects of PF4 on plasma biomarker levels following systemic polymicrobial sepsis in WT mice. (**A**) WT mice were administered CS (80 mg CS per mouse) or CS + PF4 (100 µg PF4 per mouse) for 20 min at 37ºC prior to tail vein injection. Whole blood samples were collected at 24 hr post CS challenge and subjected to proximity extension assays for quantification of selected soluble factors in platelet-poor plasma. (**B**) Platelet-poor plasma samples from (**A**) were also subjected to ELISAs for TAT and VWF as markers of coagulopathy. Mean ± SE. ^*^*P* < 0.05, ^**^*P* < 0.01, ^***^*P* <0.001, ^****^*P* < 0.0001.

Our group previously reported that PF4 reduces thrombogenicity in LPS-induced endotoxemia.^18^ To determine whether this holds true in polymicrobial sepsis, we measured thrombin-antithrombin (TAT) complexes and VWF levels in plasma from WT mice at 3, 6, and 24 hrs post-CS challenge. Both TAT and VWF levels increased in response to CS, consistent with coagulopathy. However, PF4 co-treatment reduced TAT and VWF levels at 6 hr, and thrombocytopenia 6 and 24 hr post CS challenge (**Fig. 3B**), supporting a role for PF4 in limiting sepsis-associated consumptive coagulopathy.

### PF4 enhances bacterial clearance in the pulmonary microvasculature while limiting thrombosis

We next used intravital microscopy to visualize real-time interactions between bacteria and the endothelium in the pulmonary microvasculature. WT and *PF4*^*-/-*^ mice were infused intravenously with GFP-expressing *E. coli*, and imaged for 1 hr. Upon infusion, circulating *E. coli* (green) were rapidly sequestered into the pulmonary microvasculature, followed by recruitment of platelets (red) and neutrophils (blue) (**Fig. 4A, Supplemental Videos 1-2**). *PF4*^*-/-*^ mice exhibited significantly higher *E. coli* count than WT mice 40 and 60 min post-infusion (**Fig. 4B**). *E. coli* area was initially greater in WT mice at 20 min but was significantly lower at 60 min, suggesting that endogenous PF4 facilitates early bacterial sequestration and aggregation followed by more rapid clearance at later time points (**Fig. 4B**). *PF4*^*-/-*^ mice also exhibited persistent platelet accumulation and aggregation throughout the intravital imaging period, as reflected by higher platelet fluorescent intensity and larger platelet area compared to WT (**Fig. 4C**).

**Figure 4.**
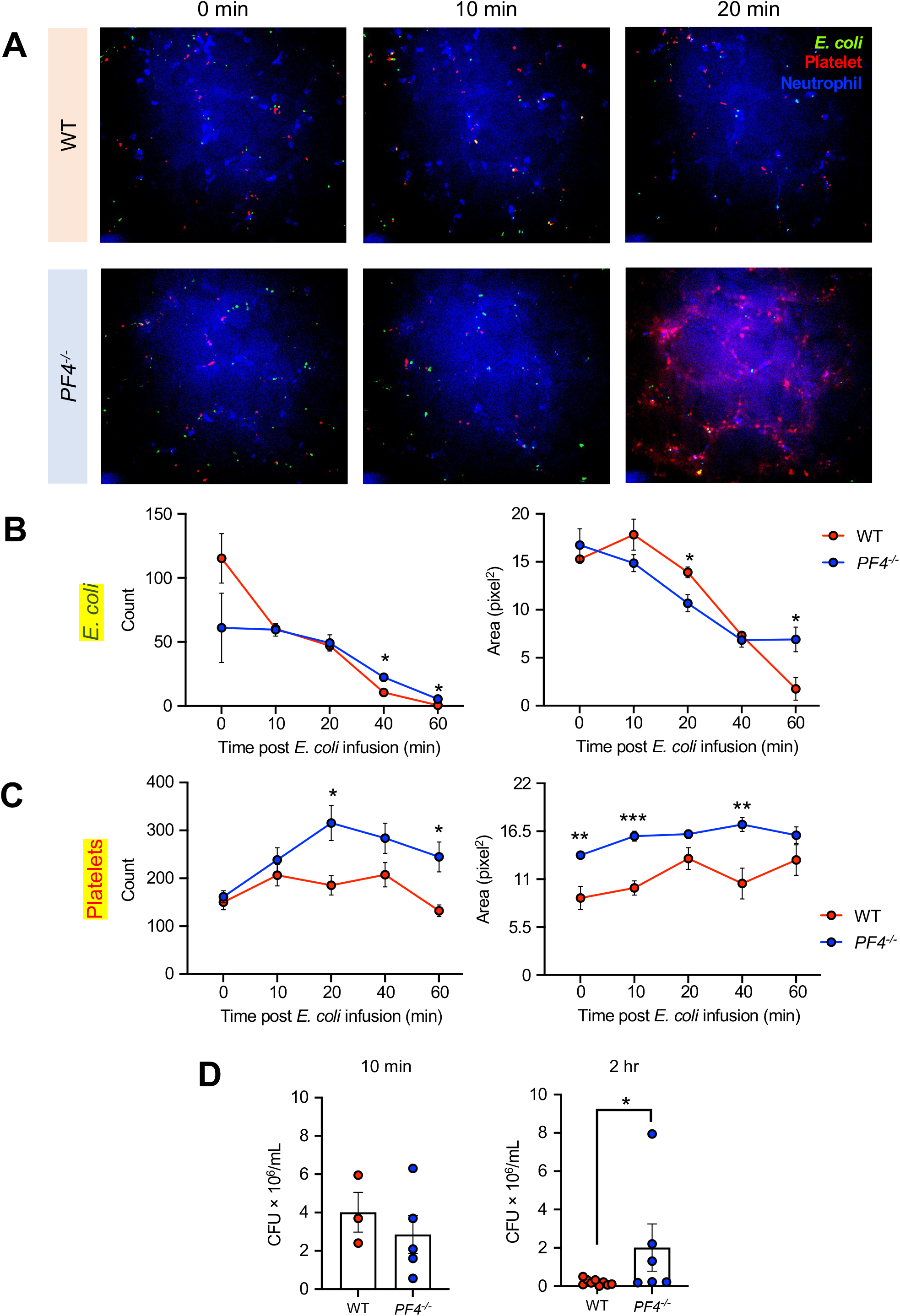
Effects of PF4 on bacterial trafficking and clearance in murine pulmonary microvasculature. (**A**) Antibodies against platelets (CD41; red) and neutrophils (Ly6G; blue) were infused intravenously into WT (top panel) and *PF4*^*-/-*^ (bottom panel) mice 20 min prior to imaging. *E. coli*-GFP (green; 1×10^9^ CFU/mouse) were infused via a jugular vein cannula, and lung images were immediately captured by confocal microscopy for 1 hr. (**B-C**) *E. coli* (**B**) and platelet (**C**) signals during intravital imaging were quantified for particle count (left) and area (right). (**D**) Lungs of mice that underwent intravital microscopy were harvested at the end of imaging experiment (2 hr post *E. coli* infusion) for CFU enumeration. A subset of mice was also administered *E. coli* (1×10^9^ CFU) through jugular vein cannula, and lungs were harvested 10 min post infusion and subjected to CFU enumeration to assess bacterial load at an early timepoint. Mean ± SE. ^*^*P* < 0.05, ^**^*P* < 0.01, ^***^*P* <0.001.

To confirm that the decrease in *E. coli* signal was due to bacterial clearance rather than photobleaching of GFP, lungs were harvested at 2 hr post *E. coli* infusion and plated for CFU enumeration (**Fig. 4D**). To ensure equal bacterial exposure at baseline, a separate cohort of WT and *PF4*^*-/-*^ mice was infused with *E. coli* and euthanized 10 min post-infusion and lung homogenates were plated for CFU analysis (**Fig. 4D**). No difference in lung CFUs was observed at 10 min, but by 2 hr, *PF4*^*-/-*^ mice exhibited significantly higher lung bacterial burden than WT mice (**Fig. 4D**), supporting that PF4 promotes bacterial clearance in the pulmonary microvasculature.

### PF4 facilitates bacterial clearance in hepatic microvasculature independent of Kupffer cells

The liver serves as a primary “firewall” for clearing blood-borne pathogens, largely through the activity of liver-resident macrophages (Kupffer cells; KC). To investigate whether PF4 influences hepatic bacterial clearance, we performed intravital imaging in WT and *PF4*^*-/-*^ mice following intravenous infusion with GFP-expressing *E. coli*. In both genotypes, *E. coli* (green) was rapidly captured by KC (blue) in the liver microcirculation. Co-localization analysis revealed no differences in KC-*E. coli* interactions between genotypes, suggesting that PF4 does not influence initial KC-mediated bacterial trapping (**Fig. 5A-B, Supplemental Videos 3-4**).

**Figure 5.**
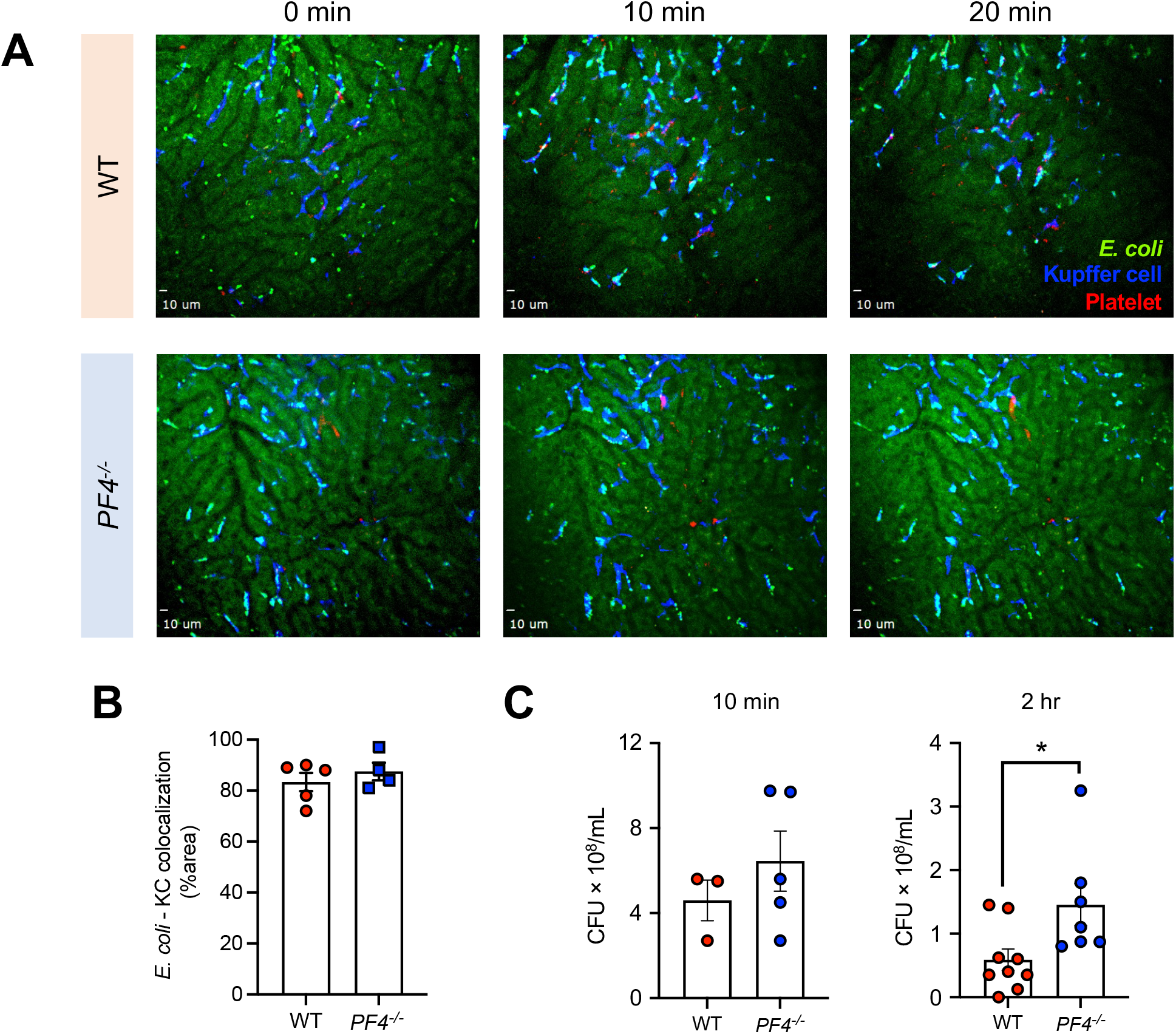
Effects of PF4 on bacterial trafficking and clearance in murine hepatic microvasculature. (**A**) Antibodies against platelets (CD41; red) and tissue-resident macrophages (Kupffer cells) (F4/80; blue) were infused intravenously into WT (top panel) and *PF4*^*-/-*^ (bottom panel) mice 20 min prior to imaging. *E. coli*-GFP (green; 1×10^9^ CFU/mouse) were infused via a jugular vein cannula, and liver images were immediately captured by confocal microscopy for 1 hr. (**B**) *E. coli* colocalization with Kupffer cells were determined. (**C**) Livers of mice that underwent intravital microscopy were harvested at the end of imaging experiment (2 hr post *E. coli* infusion) for CFU enumeration. A subset of mice was also administered *E. coli* (1×10^9^ CFU) through jugular vein cannula, and livers were harvested 10 min post infusion and subjected to CFU enumeration to assess bacterial load at an early timepoint. Mean ± SE. ^*^*P* < 0.05.

To assess bacterial burden over time, separate cohorts of WT and *PF4*^*-/-*^ mice were euthanized at 10 min and 2 hr post-*E. coli* infusion for liver CFU quantification. At 10 min, there was no difference in liver CFUs between genotypes, indicating comparable initial bacterial exposure. However, by 2 hr, *PF4*^*-/-*^ mice exhibited significantly higher liver CFUs than WT mice (**Fig. 5C**), suggesting impaired bacterial clearance in the absence of PF4.

To determine whether PF4-mediated bacterial clearance occurs independently of KCs, we depleted liver KCs in WT and *PF4*^*-/-*^ mice using clodronate liposomes.^34^ In KC-depleted animals, intravital imaging revealed that 10 min following infusion, *E. coli* formed large clusters in the hepatic sinusoids of *PF4*^*-/-*^ mice. These aggregates remained larger than those observed in WT mice over the imaging period (**Fig. 6A-B, Supplemental Videos 5-6**). As seen in the lungs, *PF4*^*-/-*^ mice also had increased platelet accumulation in hepatic microvasculature, with significantly increased platelet adhesion compared to WT mice (**Fig. 6C**). These findings suggest that PF4 facilitates bacterial clearance in hepatic microvasculature through a mechanism partly independent of KC phagocytosis, consistent with a direct role for ECs in this process.

**Figure 6.**
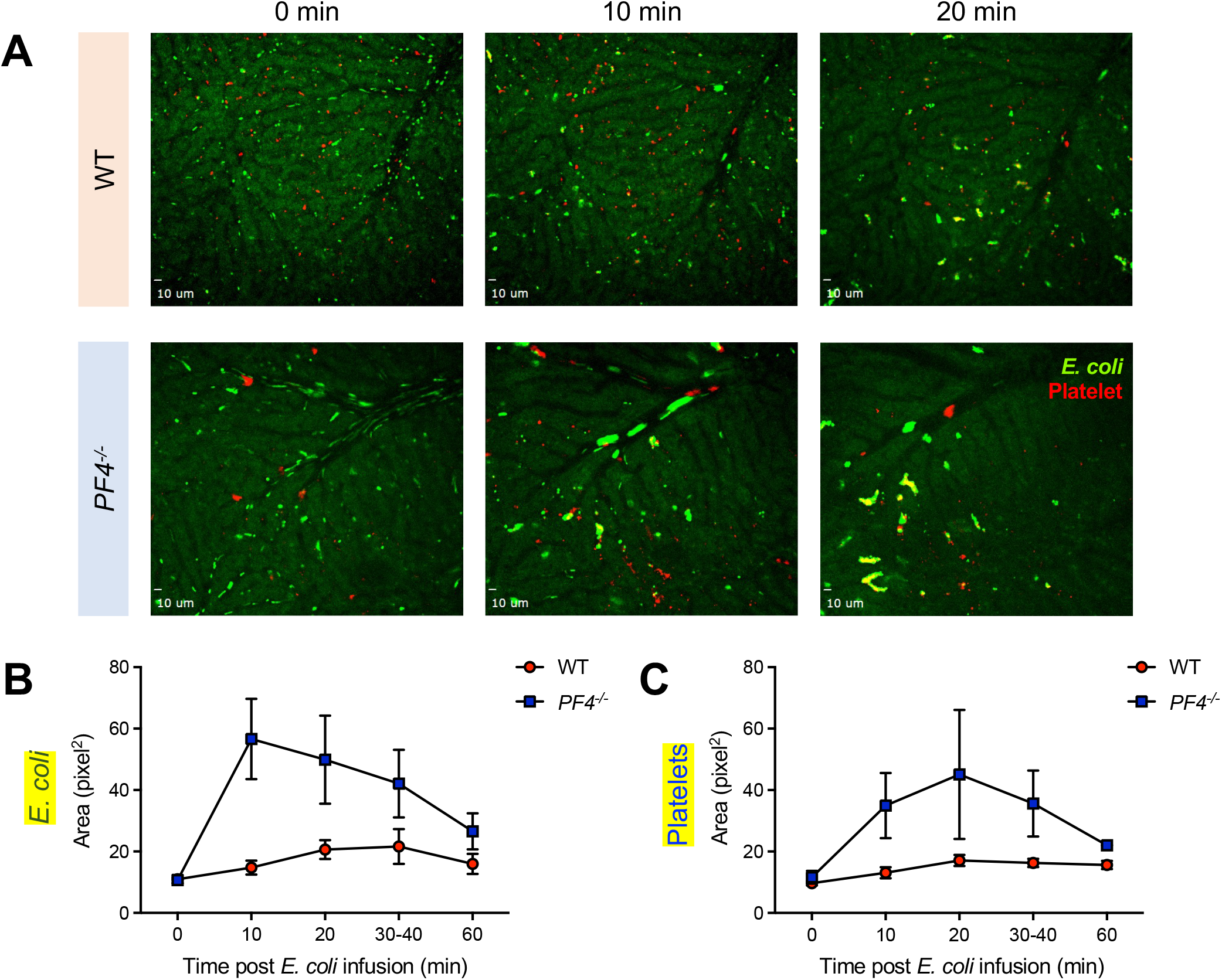
Effects of PF4 on bacterial trafficking in murine hepatic microvasculature upon Kupffer cell depletion. WT and *PF4*^*-/-*^ mice were administered clodronate liposomes to selectively deplete Kupffer cells. (**A**) Antibodies against platelets (CD41; blue) and tissue- resident macrophages (Kupffer cells) (F4/80; red) were infused intravenously into WT (top panel) and *PF4*^*-/-*^ (bottom panel) mice 20 min prior to imaging. *E. coli*-GFP (green; 1×10^9^ CFU/mouse) were infused via a jugular vein cannula, and liver images were immediately captured by confocal microscopy for 1 hr. (**B-C**) *E. coli* (**B**) and platelet (**C**) signals during intravital imaging were quantified for total area. Mean ± SE.

## DISCUSSION

This study reveals a critical role for platelet PF4 in strengthening host defense against bloodstream infections by promoting endothelial uptake and clearance of bacteria. We show that PF4 binds to bacterial surfaces and to endothelial receptors, facilitating bacterial internalization, lysosomal degradation, and systemic clearance. These effects are accompanied by reduced inflammation, coagulopathy, and mortality in murine models of sepsis. Beyond expanding the known functions of PF4, our findings support a broader concept that ECs, long recognized as passive barriers, actively participate in innate immune defense during systemic infection.

PF4 is structurally distinct from canonical CXC chemokines in that it lacks the N-terminal ELR motif, preventing activation of CXCR1 and CXCR2, but not by CXCR3B. On the other hand, its tetrameric conformation and strong cationic charge confer high affinity for polyanionic ligands, including bacterial surface components, heparan-sulfate-rich proteoglycans in the endothelial glycocalyx, nucleic acids, and VWF. These properties underlie its ability to form immune complexes, stabilize NETs, and participate in host-pathogen interactions. Our data extend these roles by showing that PF4 binds directly to bacteria to promote their uptake by ECs via both CXCR3B engagement and GAG-mediated interactions. At lower concentrations, PF4 enhances bacterial endocytosis, whereas higher concentrations appear to inhibit uptake, suggesting a threshold effect. This may be influenced by bacterial aggregate size and surface repulsion. Prior work has shown that bacterial aggregate size significantly affects phagocytosis by neutrophils, with efficient uptake of aggregates ≤ 5μm in diameter but impaired phagocytosis of larger bacterial clusters.^35^ Similarly, excess PF4 may saturate bacterial and endothelial surfaces, causing electrostatic repulsion. Dose-dependent modulation of bacterial clearance may have implications for therapeutic PF4 use *in vivo*.

Our findings also contribute to a growing appreciation of ECs as immune effectors. While ECs are well known to express pattern recognition receptors and inflammatory mediators, accumulating evidence suggests that they can perform macrophage-like functions, including phagocytosis and bacterial killing. For example, primary human umbilical vein endothelial cells (HUVECs) internalize *Listeria monocytogenes* via a formin-dependent, phagocytosis-like process and traffic them to lysosomes for degradation, demonstrating endothelial lysosomal activity in vivo during infection.^24^ Similarly, liver sinusoidal ECs actively clear and degrade circulating particles like bacteriophages via lysosomal pathways in situ.^24^ We show that PF4 promotes uptake of bacteria by ECs, and enhances trafficking to lysosomes, potentiating intracellular killing—all without compromising membrane integrity or inducing generalized EC activation, as assessed by VWF and LDH release. These antimicrobial functions appear particularly important in the pulmonary and hepatic microvasculature, where ECs represent the first line of defense against circulating pathogens. In the absence of Kupffer cells, PF4 was sufficient to enhance hepatic bacterial clearance, supporting a direct role for ECs in pathogen removal.

PF4 also modulates the thromboinflammatory milieu of sepsis. Prior work has shown that PF4 can bind to NET DNA to attenuate NET-induced thrombosis.^8,18^ In the present study, PF4 administration reduced plasma levels of TAT complexes and VWF in septic mice, consistent with reduced coagulation activation and endothelial stress. In parallel, PF4-deficient mice exhibited excessive platelet aggregation in the lung and liver. These findings suggest that endogenous PF4 helps limit uncontrolled platelet activation, possibly by reducing EC activation. This contrasts with earlier reports that PF4 promotes platelet aggregation in response to sterile stimuli,^6^ pointing to a potentially distinct role for PF4 in infectious contexts. Collectively, these data align with the concept of “immunothrombosis,” in which coagulation is harnessed for host defense, but must be tightly regulated to avoid tissue injury and organ failure.^36,37^ Although our data highlight a protective role for PF4 in infection, they may also relate to recent work showing that PF4 treatment in mice reduces age-associated inflammatory changes, including elevated type I interferon signaling and neutrophil activation, to decrease neuroinflammation and improved cognition.^38^

We have considered the translational implications of our findings. Patients with thrombocytopenia, platelet dysfunction, or platelet hyper-reactivity and ⍰-granule storage pool exhaustion, such as those with trauma, cancer, or sickle cell disease, may be at increased risk for sepsis-associated organ dysfunction due to impaired PF4-mediated bacterial clearance.^39^ Replacing PF4, or using agents that mimic its opsonic functions, could represent a novel host-directed strategy to enhance bacterial clearance while dampening inflammation and thrombosis. Although no adverse effects were reported in a small study of individuals infused with PF4 as a heparin reversal agent,^40^ treatment with exogenous PF4 requires caution. Circulating PF4 rapidly binds to the endothelial glycocalyx, plasma levels typically underestimate true vascular stores.^41^ Therefore, patients with platelet hyper-reactivity, even in the setting of thrombocytopenia, may already have optimal levels of vascular PF4, and additional PF4 could impair bacterial clearance. Furthermore, PF4 tetramers can bind to heparin and other polyanions to form complexes that expose neoepitopes recognized by pathogenic anti-PF4 antibodies. This can trigger platelet activation and thrombosis, as seen in thrombotic anti-PF4 immune disorders such as heparin-induced thrombocytopenia (HIT), vaccine-induced thrombocytopenia and thrombosis (VITT), and related conditions.^29,42,43^ One strategy explored by our group is the use of a non-immunogenic antibody that stabilizes PF4-polyanion complexes, enhancing the effects of endogenous PF4 without exposing patients to supraphysiologic PF4 levels.^8^ Optimizing dose, delivery method, and modifications to reduce immunogenicity will be essential in therapeutic development.

In summary, our findings establish PF4 as a key mediator linking platelets, endothelial cells, and innate immune defense. PF4 opsonizes bacteria, promoting their uptake and clearance by ECs, while simultaneously limiting systemic inflammation and thrombosis. These dual antimicrobial and immunomodulatory functions suggest that PF4 operates at the nexus of hemostasis and immunity. Notably, because PF4 facilitates bacterial clearance through host-directed mechanisms rather than direct antimicrobial activity, its protective effects may extend to infections caused by antibiotic-resistant pathogens. Harnessing endogenous defense pathways like PF4 may represent a promising strategy to improve patient outcomes in sepsis and multidrug-resistant infections.

### Study ethics approval

Animal procedures were approved by the Institutional Animal Care and Use Committee (IACUC) in accordance with the NIH Guide for the Care and Use of Laboratory Animals (National Academies Press, 2011) and the Animal Welfare Act.

## Acknowledgments

This work was supported by a National Institutes of Health National Heart Lung and Blood Institute grant R00HL177830 (K.G), and an American Society of Hematology Scholar Award (A.T.P.N.)

## Authorship

A.T.P.N. conducted experiments, analyzed data, and prepared the initial and revised drafts of the manuscript. W.O. performed hepatic intravital imaging studies, analyzed data, and provided research guidance. A.S. carried out polymicrobial sepsis studies. H.K. performed pulmonary intravital imaging studies and data analysis. J.O. purified PF4, provided research guidance, and assisted with murine sepsis studies. A.S. assisted with endothelial cell culture and overall guidance. V.B. and N.L. developed the murine sepsis model and maintained the mouse colony. L.R. provided overall guidance related to intravital microscopy and data interpretation. I.C. performed SEM studies. Z.M. and C.D. performed Olink proteomic studies. M.G. shared GFP-*E. coli* and provided research input. V.N. contributed to microbial studies and manuscript preparation. M.P. provided research guidance, expertise related to PF4 biology, and assistance with manuscript preparation. K.G directed the project, interpreted data, and oversaw manuscript preparation.

